# Quantifying Variability in Attentional Shifts During Target Switching Using Steady-State Visual Evoked Potentials

**DOI:** 10.1101/2023.10.21.563393

**Authors:** Moran Eidelman-Rothman, Omer Reuveni, Andreas Keil, Lior Kritzman, Dominik Freche, Hadas Okon-Singer, Nava Levit-Binnun

## Abstract

A large body of ssVEP research has provided significant insights into the temporal dynamics of attentional processes. While these studies focused primarily on group level inspection, there is a need for further research employing methodological approaches that enable the examination of individual-level variability. This is particularly relevant in psychological contexts, where such measures are linked to various cognitive and clinical outcomes. In the present study, we aimed to measure and discern attentional shift processes, examining both group and individual subject dynamics. We utilized EEG frequency tagging to examine attentional engagement, disengagement, and reengagement while participants switched focus between four flickering stimuli. Analysis of ssVEPs revealed significant changes in amplitude between attentional conditions. Specifically, group-level results indicated an increase in activity during engagement with the first target, followed by a decrease upon disengagement, while reengagement with the second target showed a corresponding increase in activity occurring on average 110 ms prior to disengagement. Distinct individual patterns emerged, with participants showing either disengagement, reengagement, both, or no shifts. Notably, the timing and order of these attentional shifts varied considerably across individuals relative to target switch events. These findings demonstrate the ability of this approach to quantify attentional shifts on both group and individual levels, providing a foundation for further research into individual differences in attentional control, which may have implications for understanding adaptive and maladaptive psychological functioning.

**Highlights:**

- EEG frequency tagging captures attentional engagement, disengagement, and reengagement processes.
- Group-level ssVEP analysis reveals that attentional reengagement occurs 110 ms before attentional disengagement.
- Individual-level analysis demonstrates diverse patterns of attentional shift timing and order.
- The experimental framework successfully quantifies attentional dynamics on the group and individual levels.

## Introduction

Moment by moment our senses are exposed to continuous information from the external environment as well as our internal world, such as thoughts and bodily sensations. Attention processes allow us to deal with the limited capacity of perceptual and cognitive systems that use this information by prioritizing certain types of information and suppressing others (Desimone & Duncan, 1995). Consequently, attention modulates many essential cognitive processes such as perception, working memory and executive functions (Luck & Gold, 2008).

A key aspect of attentional processes is the ability to execute attentional shifts—transitions between inputs that include engagement with relevant stimuli, disengagement from irrelevant inputs, and reengagement with new, more relevant ones (Petersen & Posner, 2012; Posner, 1980). These shifts are essential for flexibly managing cognitive resources in an efficient way, enabling perceptual accuracy, decision-making, and adaptability in complex environments where the relevance of stimuli frequently changes (Carrasco, 2011; Luck et al., 2021; Lutz et al., 2008; Theeuwes, 2014). Inefficiencies in managing these shifts are frequently reflected in altered temporal dynamics of these processes (e.g., delays in disengaging attention) and serve as critical markers of cognitive effort and attentional control (Eysenck et al., 2007; Luck & Gold, 2008). Flexibly managing attentional resources is vital not only for cognitive efficiency but also for maintaining psychological resilience in the face of fluctuating demands (Dahl et al., 2020; Lutz et al., 2015). Therefore, capturing these patterns with precision is crucial for understanding the mechanisms underlying attentional regulation and its role in adaptive and maladaptive psychological functioning (Eysenck et al., 2022, 2022; Lutz et al., 2015; MacLeod et al., 2019).

Maladaptive psychological conditions such as anxiety and depression are frequently characterized by inefficiencies in attentional shifts rather than an inability to shift attention (Eysenck et al., 2022; Koster et al., 2017). For example, individuals with anxiety often exhibit attentional shift biases that lead to fast and more dominant engagement with negative stimuli or difficulty disengaging from them, resulting in altered reaction times. These inefficiencies are not confined to emotional contexts but are also evident in non-emotional tasks. For instance, in antisaccade paradigms, anxious individuals commonly show prolonged latencies in disengaging from irrelevant inputs and redirecting attention toward task-relevant stimuli, even in neutral settings (Berggren & Derakshan, 2013; De Lissnyder et al., 2011). Similarly, ERP studies of non-emotional attentional tasks across conditions such as anxiety, depression, and ADHD reveal variations in the latencies of neural components which are linked to attentional shifts (Ahumada-Méndez et al., 2022; Hepsomali et al., 2019; Hsieh et al., 2021). These inefficiencies, which reflect increased effort and altered timing, are thought to play a central role in the etiology, persistence, and exacerbation of symptoms (Abado et al., 2020; Bar-Haim et al., 2007; Derakshan, 2020; Dolcos et al., 2020; Okon-Singer, 2018). Therefore, examining the temporal dynamics of attentional shifts, particularly their timing and effort, provides critical insights into both adaptive and maladaptive functioning.

One of the most prominent methods used in recent years to gain more insight into the time course and temporal dynamics of attentional shifts is the steady-state visual evoked potentials (ssVEPs). SsVEPs are oscillatory cortical responses evoked by the periodic modulation of a sensory stimulus over a period of several seconds. These responses are characterized by rhythmic engagement of groups of neurons that are responsive to the stimulus and can be identified as oscillatory brain responses matching the frequency of the stimulus (Vialatte et al., 2010). This tool is especially useful in attentional research, with the time-varying amplitude of ssVEPs providing a continuous measure of attentional resource allocation. Specifically, increased ssVEP amplitudes are an indication of heightened attention directed towards locations, features and/or objects while decreased ssVEP amplitudes reflect decrease of attention towards such inputs (Andersen et al., 2011; Andersen & Müller, 2010; De Echegaray et al., 2024; Kritzman et al., 2022; Norcia et al., 2015; Wieser et al., 2016).

SsVEPs provide several advantages when studying attention allocation related processes. These include high signal to noise ratio (SNR) due to the narrow band filtering of specific frequencies and the possibility of following attentional processes over longer periods of time without being constrained to time locked data. Moreover, ssVEPs enable the measurement of attentional allocation towards a specific stimulus even when it is embedded in a complex scene using the frequency tagging technique. In this technique the response to various stimuli can be isolated by periodically modulating the luminance of the stimuli (i.e., flickering) at distinct frequencies. (Norcia et al., 2015; Wieser et al., 2016). For example when presenting two embedded random dot kinematograms (RDKs) of different colors flickering in different frequencies in the same spatial location, the mere direction of covert attention to one of the two RDKs (i.e., to a specific color) results in enhanced amplitude of the time-varying signal in the attended frequency (Adamian et al., 2019; Andersen et al., 2011; Andersen & Müller, 2010; Forschack et al., 2017; Gundlach et al., 2022).

Studies utilizing this approach have also provided significant insights into the time-sensitive nature of attentional shifts. For example, utilizing a design in which participants had to attend to either one of two superimposed RDKs, Andersen and Müller, (2010) found that, amplification of ssVEP amplitude in response to the attended stimulus occurred prior to the reduction in the amplitude associated with the stimulus that participants needed to ignore. This finding that indicated that engagement to the relevant stimulus occurs prior to the suppression of the irrelevant one underscores the independence of attentional processes. Later studies (Müller et al., 2016; Vieweg & Müller, 2021) further expanded these insights, showing that reengagement with a new target begins before full disengagement from a previous one. Moreover, the latency of this switch was longer for attentional shifts that occur across different features (e.g., color to orientation) as compared with attentional shifts within the same features (e.g., one color to a different color). These studies highlight the ability to use ssVEPs as measures of disengagement and reengagement of attention switching and the use of latency of the shift as indicative of increased cognitive demands and attentional effort (Eysenck et al., 2007). By capturing these latency differences, ssVEPs provide a precise measure of temporal dynamics.

Importantly, the high signal-to-noise ratio (SNR) of ssVEP offers a significant advantage—not only for studying attentional processes at the group level but also for examining them on an individual subject basis (e.g., Gundlach et al., 2020; Nunez et al., 2015). While group-level analyses provide valuable insights into general patterns of attentional dynamics, they often mask the variability in attentional processes between individuals (Petersen & Posner, 2012). Understanding these individual differences is crucial, as they can reveal distinct attentional profiles that have important implications for both adaptive and maladaptive cognitive functioning. For example, while anxiety-linked attentional biases are reliably observed at the group level, individuals may not exhibit stable or consistent attentional patterns across contexts (MacLeod et al., 2019; Zvielli et al., 2015). Several attempts have succeeded to demonstrate the ability of ssVEPs to capture individual differences in the context of attention engagement and suppression. For instance, Gundlach et al. (2022) demonstrated that participants with the greatest ssVEP amplitude facilitation towards an attended stimulus did not necessarily show the highest suppression towards the unattended stimulus. This finding not only underscores the independence of attentional engagement and suppression processes, but also highlights the importance of examining individual differences in these mechanisms, which may reveal distinct attentional profiles not captured by group-level analyses.

Building upon previous work demonstrating the utility of ssVEPs in capturing the temporal dynamics of attentional switching and identifying individual attentional profiles, the present study aimed to refine and expand the ability to measure and discern attentional engagement, disengagement, and reengagement processes, alongside their latencies, at both the group and individual levels. To achieve this, we integrated well-established methods into a novel approach designed to examine these dynamics comprehensively. Using the frequency tagging technique, participants were required to switch their attentional focus between stimuli presented simultaneously in the same visual space. We hypothesized that ssVEP responses would reflect distinct patterns of neural activity corresponding to attentional shifts, with increases in amplitude during engagement, decreases during disengagement, and subsequent increases during reengagement. Given the known variability in attentional processes, we further hypothesized significant individual differences in latency, reflecting variations in attentional shifts across participants. Ultimately, this approach lays the groundwork for future research aimed at assessing individual differences in attentional control and supporting the development of interventions to enhance these processes.

## Method

### Participants

Based on previous studies with similar designs (Gundlach et al., 2022; Vieweg & Müller, 2021) we aimed to reach a final sample of thirty participants. Forty-one participants were eventually recruited in ages between 18 - 35 with normal color perception from the Reichman university’s psychology department for course credits. Exclusion criteria included current or past neurological or psychiatric disorders. Twelve participants were excluded from analysis: Six due to equipment malfunction, one due to low task performance (for details see “Behavioral accuracy” in the Results section), and five due to excessive lateral eye movements (for details see “Data preprocessing” in the Methods section). Thus, the final experimental sample included 29 participants (18 females; 24 right-handed, mean age = 24.48 (SD = 3.26). The study was approved by the university’s ethics committee.

### Experimental Stimuli

In line with previous designs (e.g., Andersen et al., 2011a), two overlapping random dot kinematograms (RDKs) of different colors (red, RGB = 1, 0, 0, and green, RGB = 0, 0.8, 0) were presented on both the left and the right side of a central fixation arrow (see Figure 1). Each of the total four RDKs consisted of a population of 700 dots flickering at either 8, 10, 12 or 15 Hz. For each participant, the flickering frequencies were pseudo-randomly assigned to each of the four RDKs. The outer edges of each circle shaped RDK had a visual angle of 13.92° and the circles’ inner edges were located 0.92° from a fixation point. Each dot occupied 0.15° and was moving randomly to one of four cardinal directions. Coherent motion events in the RDK were created for the behavioral task (see section Experimental task) by including short periods (333 ms) during which 80 percent of the dots in the attended RDK moved in the same direction. Stimuli were presented against a black background on a 24-inch monitor display (AOC G2460PF), with resolution of 1920 X 1080, and a refresh rate of 60 frames per second.

**Figure 1.**
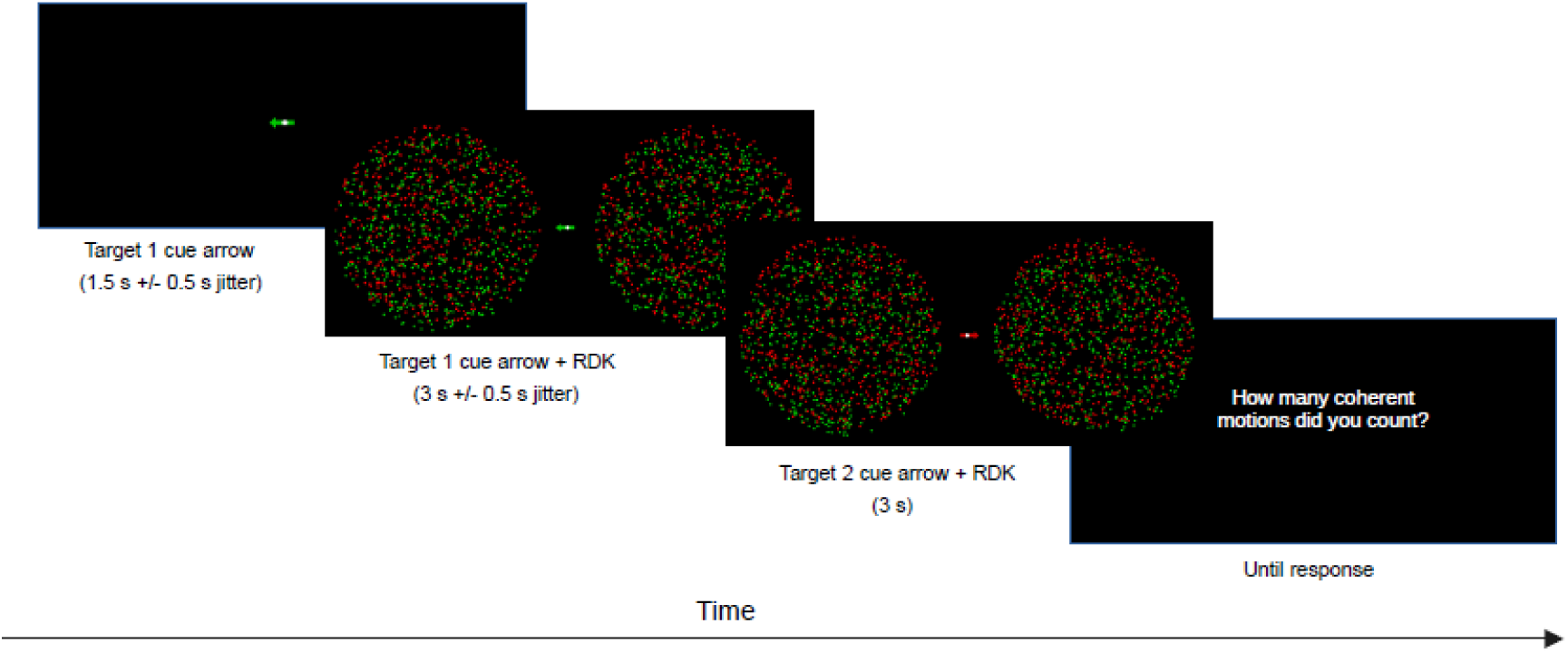
The experimental trial design and configuration (not to scale). Each trial started with 1.5 s ± 0.5 s presentation of a red or green cue arrow, pointing either to the left direction, the right direction or to both directions, followed by the appearance of four random dot kinematograms (RDKs; four clouds of dots, each flickering in one of the stimulation frequencies). After a random interval of 2.5 to 3.5 s the cue arrow changed its color, or its direction or both (“target switch”), and all flickering dots remained on the screen until a 6 s interval elapsed. Participants had to direct their attention to the target according to the cue arrow. At the end of each trial participants were asked to report the number of coherent motions they counted (0,1 or 2).

### Experimental Task

Each trial started with the presentation of a red or green cue arrow, pointed either to the left direction, the right direction or a bidirectional arrow pointing to both directions, as shown in Figure1. Thus, there were overall 6 possible attentional targets which occurred with equal probability: red-right, red-left, red-bidirectional, green-right, green-left, and green- bidirectional. A white fixation dot appeared at the center of each arrow during the entire task. Following a random interval of 1 to 2 s (average of 1.5 s across trials ± 0.5 s jitter), all four RDKs appeared on the screen. To obtain measures of attentional disengagement and reengagement (as defined in the introduction), a target shift occurred in the middle of the trial: after a random interval of 2.5 to 3.5 s (average of 3 s across trials ± 0.5 s jitter, to avoid temporal expectancy effects) the cue arrow changed either its color, its direction or both (“target switch”), and all flickering dots remained on the screen until a 6 s interval elapsed. In each trial, the participants were given the task of attending to a specific RDK (attended target) specified by the first cue arrow, and to switch their attention to a new target after the arrow has changed. In addition, to assure that participants were engaged with the task they were asked to detect short intervals of coherent motions in the attended stimulus, which occurred in 30 % of the trials. Coherent motion events could appear in the first attended target (before the condition switch), in the second attended target (after the switch) or in both, and their onset did not occur earlier than 0.4 s following the onset of the RDKs or the target switch and not later than 0.25 s before the switch or the end of the trial. At the end of each trial, participants were asked to report the number of coherent motions they detected using a button press (possible answers were 0, 1 or 2 coherent motions). In this design, we aimed to measure covert attention to avoid motor-related confounds, which could influence attentional processing. While participants were asked to detect coherent motion, no immediate behavioral response was required during the trial. However, behavioral accuracy was assessed at the end of each trial, where participants reported the number of coherent motion events they detected. This approach allowed us to ensure that participants remained engaged and accurately directed their attention to the target stimuli. The experiment included a total of 180 trials, presented in 6 blocks of 30 trials with one break between the blocks (total of 5 breaks). Participants were instructed to maintain their gaze on the white fixation dot located at the center of the cue arrow during the entire task.

### Procedure

Upon arrival at the lab, participants were given an overview of the experiment, and after signing a consent form, they were connected to the EEG set up. The experiment took place in a dimly lit room. Participants were seated approximately 0.65 meters from the screen.

Following a 6-min baseline recording (3 minutes resting state with eyes open and 3 minutes with eyes closed) they were provided with instructions for the main task and performed a training session of 18 trials or more, until the experimenter confirmed that participants understood the task (via conversation and observation of their performance during the practice trials). They then performed the main task followed by an additional resting state recording performed as described above.

### Data Acquisition

Continuous electroencephalographic (EEG) activity was recorded using the ActiveTwo BioSemi system (BioSemi, Amsterdam, The Netherlands). Recordings were obtained from 64 scalp electrodes based on the ten-twenty system, as well as from two electrodes placed on the left and right mastoids. The electrooculogram (EOG) generated from blinks and eye movements was recorded from four facial electrodes: two approximately 1 cm above and below the participant’s right eye, one approximately 1 cm to the left of the left eye, and one approximately 1 cm to the right of the right eye. As per BioSemi’s design, the ground electrode during acquisition was formed by the Common Mode Sense active electrode and the Driven Right Leg passive electrode. All bioelectric signals were digitized on a laboratory computer using ActiView software (BioSemi). Data was digitized at a sampling rate of 2048 Hz, utilizing a high pass filter set at 0.16 Hz and a low-pass online filter set at 100 Hz. The electrode offset was kept within ± 25 mV according to the user manual. In order to achieve greater synchronization between visual stimuli presentation and EEG recordings, we used a set of photodiodes that sent the stimulus triggers with high precision as they appeared on the screen (Peterson et al., 2014).

### Behavioral Accuracy

Behavioral accuracy scores in the coherent motion detection task were calculated using the following formula which was based on Pollatos & Schandry, (2004).

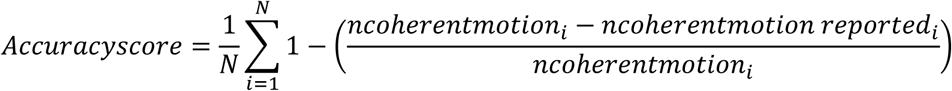

Where N is the total number of trials in which coherent motions were presented and n is the number of coherent motions in a specific trial (i).

### Data Pre-processing

Offline pre-processing was performed in Matlab (The MathWorks; R2020a), using in-house built scripts, based on functions from EEGLAB (Delorme & Makeig, 2004) and Fieldtrip (Oostenveld et al., 2010). The continuous data were bandpass filtered offline in the 1- 60 Hz range (Hamming windowed sinc FIR filter) and were then down-sampled to 512 Hz.

Subsequently, the data was segmented into 8 s epochs from 1 s before the onset of the RDKs to 1s after the offset of the RDKs. The data was then re-referenced to the average of both mastoids. Blinks and eye movement artifacts were identified and corrected utilizing the Second Order Blind Identification method (Belouchrani et al., 1997) and the IClabel (Pion- Tonachini et al., 2019). In order to assure that amplification/attenuation of response to the different attentional targets can be attributed to the allocation of attention and is not the result of directing gaze towards and away from the attended targets, participants with excessive lateral eye movements were excluded from the analysis (five participants). Further rejection of artifactual trials and interpolation of noisy electrodes were performed using a Statistical Control of Artifacts in Dense Arrays Studies (SCADS) method, fully described in (Junghöfer et al., 2000).

### SsVEP Analysis

To assure that selective attention was solely based on the stimulus color and direction, with no interference from coherent motion events, trials containing coherent motions were excluded from the analysis. The current analysis was focused on attentional allocation towards unidirectional stimuli only (i.e., attending to RDKs on either the left or the right side of the fixation arrows), such that trials containing bidirectional stimuli (i.e., trials in which subjects needed to attend simultaneously to RDKs flickering on both sides of the fixation arrows) were not included. These conditions were included in the experiment to assess attentional processes related to the narrowing and widening of attention, which is beyond the scope of the current paper.

### SsVEP Signal Quality Evaluation

To visualize and evaluate the quality of the ssVEP signal, the pre-processed time-domain EEG data was averaged across trials, posterior electrodes (‘OZ’, ‘POZ’, ‘O1’, ‘O2’), conditions and subjects. In addition, the frequency content of the data was estimated by subjecting the trial-averaged time domain data to a fast Fourier transform (FFT) analysis to verify that the flickering dots reliably evoked steady-state responses in the expected stimulation frequencies. This analysis was conducted on the data within a time window extending from one second post RDK onset to one second pre RDK offset.

### Rhythmic Entrainment Source Separation (RESS) Analysis

SsVEP responses were extracted via rhythmic entrainment source separation (RESS; Cohen & Gulbinaite, 2017). In this analysis, a map of spatial weights across all electrodes is created in order to extract ssVEP signal, without relying on a priori or post-hoc electrode selection. Specifically, in RESS analysis linear spatial filters are created to maximally differentiate the covariance between signal in the flicker frequency and neighboring frequencies, thereby the SNR at a particular frequency is optimized, for each participant. After obtaining signal and neighboring covariance matrices, a generalized eigen decomposition is computed, and the eigenvector with the largest eigenvalue is used as channel weights to obtain a single component time course, which reduces multiple comparisons across channels in statistical testing. RESS was performed on the entire pool of artifact-clean trials, including trials that contain bidirectional targets that were otherwise not analyzed (see section SsVEP analysis). Interpolated electrodes (see section Data pre-processing) were not included in the analysis (Cohen & Gulbinaite, 2017). A single RESS filter was created for the four distinct stimulation frequencies by computing separately four covariance matrices and then averaging them together, to yield a single signal covariance matrix (Cohen & Gulbinaite, 2017). A single reference (“noise”) covariance matrix was constructed similarly. A generalized eigen decomposition was then computed on the signal and reference covariance matrices as described above. The spatial filter (i.e., weights) was then applied, by multiplying it by the EEG electrode time series.

### The Time-varying ssVEP Amplitude

For each stimulation frequency, we extracted the full length of all trials in which this frequency was the attended target before the switch (i.e., target 1). Similarly, for each stimulation frequency, trials in which this frequency was the attended target after the switch (i.e., target 2) were extracted (see Figure 2). Thus, two attentional conditions were formed: 1) “engage target 1 - disengage target 1” and 2) “ignore target 2 - reengage target 2”.

**Figure 2.**
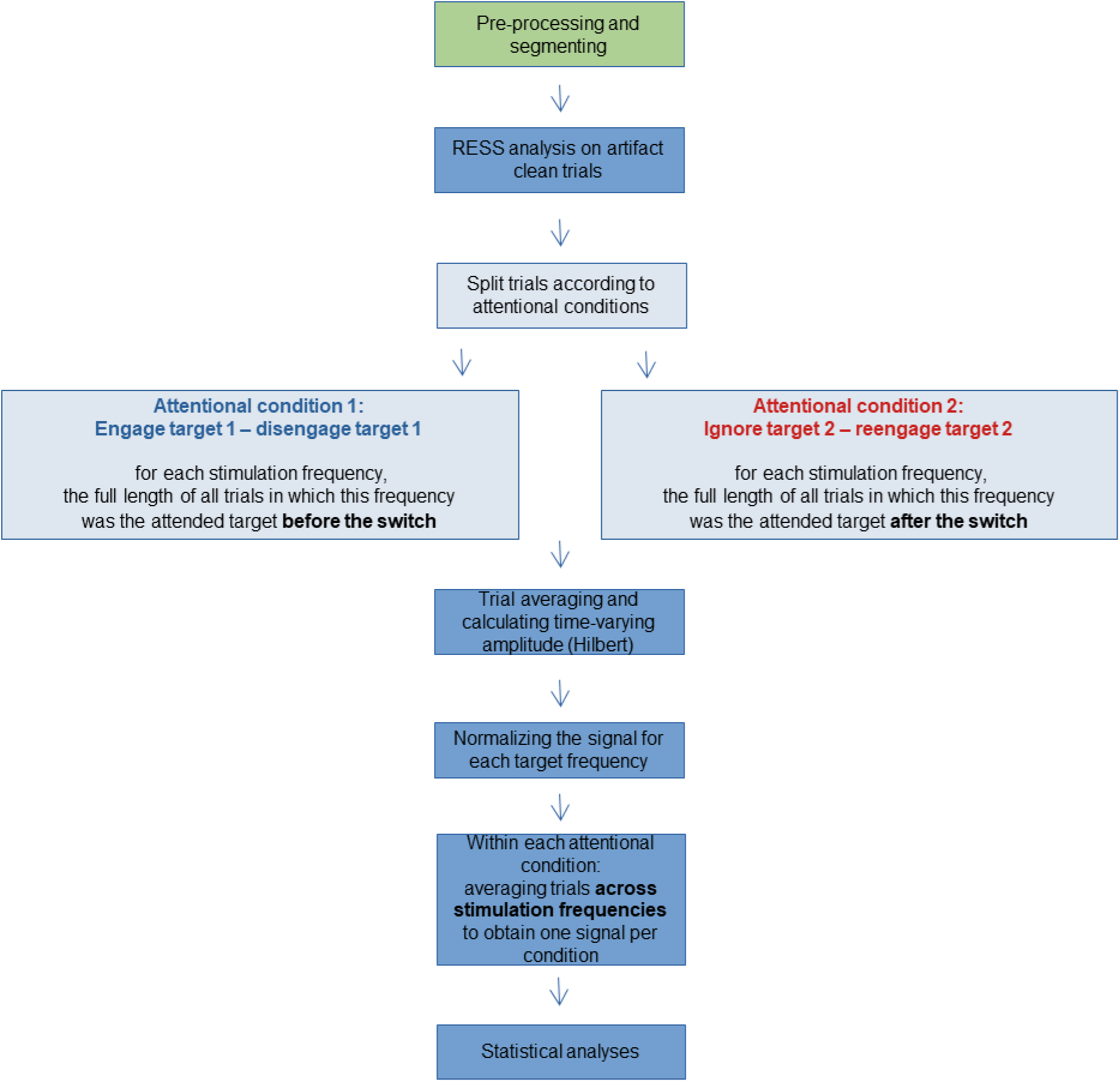
Main steps in data analysis. Following the initial steps of data preprocessing and segmentation, the entire pool of artifact-clean trials was subjected to RESS analysis, which allows for the extraction of ssVEP signals without relying on a priori or post-hoc electrode selection. The RESS procedure resulted in a single time series per trial, for each participant. The trials were then split according to the two attentional conditions defined: 1) “engage target 1 - disengage target 1” and 2) “ignore target 2 - reengage target 2”. Next, for each attentional condition, the trials of each stimulation frequency were averaged separately and subjected to Hilbert transform, to calculate the time-varying ssVEP amplitudes. The data was then normalized, which enabled the averaging of trials across the different stimulation frequencies. The normalized data were averaged within each attentional condition, resulting in a single signal per attentional condition, which was then used for statistical analysis.

For each stimulation frequency the time-varying amplitude envelopes in each of these attentional conditions were extracted by subjecting the RESS time series to Hilbert transform, as described in FreqTag toolbox (Figueira et al., 2022; using the function freqtag HILB). In the first step, trials for each frequency and each attentional condition were averaged separately. Next, each averaged signal was baseline corrected by subtracting the mean amplitude of a 0.5 s pre-stimulus period (i.e., prior to RDK onset), from each time point along the trial. The data from each frequency was then narrow band filtered (in the stimulation frequency range ± 0.5; 9th order Butterworth). The amplitude was extracted as the absolute value of the Hilbert transformed analytic signal (see Figure 2 for the full procedure). To maintain the temporal resolution while reducing noise, we used a moving average with approximately 10% of the data as the window size to smooth the Hilbert transform product of the ssVEP data.

To be able to average trials within the same attentional condition across different stimulation frequencies, the data was normalized to account for the typical ‘1/*f*’ distribution of power spectrum data, (i.e., the decrease of amplitude (A) with increasing frequency), (Andersen et al., 2011; Gyurkovics et al., 2021). As dictated by the experimental design, all four stimulation frequencies were presented in each trial, and trials differed from each other by the frequency at which the attended target was flickered. Therefore, the normalization was performed, as follows (as described in the equation below): For each participant (i), for each attentional condition (k: “engage target 1 - disengage target 1”, “ignore target 2 - reengage target 2”) and each of the stimulation frequencies (j: 8 Hz, 10 Hz, 12 Hz, 15 Hz), the amplitude of the attended frequency was divided by the weighted mean of that same frequency’s amplitude when derived from all possible stimulation conditions (nT = number of trials in the specified frequency condition). For example, the normalized amplitude of 8 Hz target frequency was calculated by dividing its amplitude in the attend 8 Hz conditions, by the weighted mean of the 8 Hz amplitude measured in all possible target frequency conditions (i.e., when the attended target was either 8 Hz, 10 Hz, 12 Hz, 15 Hz).

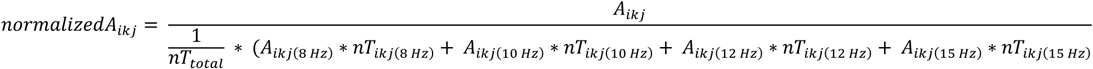

Thus, a normalized amplitude value > 1 signifies a greater response to the attended target (e.g., red-left) over the response to all possible flickering stimuli (e.g., red-left, green-left, red-right, green-right), whereas a normalized amplitude value < 1 signifies a decreased response to the attended target (e.g., red-left) over the response to all possible flickering stimuli (e.g., red-left, green-left, red-right, green-right; Figure 2). Following normalization, the signals were averaged across target frequencies in each attentional condition for each participant, to yield a single time series per attentional condition, which were submitted to statistical analysis. The main steps of the analysis are summarized in Figure 2.

### Statistical Analysis

For statistical analysis, two time windows were defined per each attentional condition: an early time window of 2 s before the target switch starting 0.5 s after the onset of the RDKs, and a later 2 s window starting 0.5 s after the target switch. These intervals were chosen in order to avoid contaminating responses related to RDK onset and to the target switch cue. This resulted in a 2×2 within-subject design of two time windows (before switch, after switch) by two attentional conditions (“engage target 1 - disengage target 1”, “ignore target 2 - reengage target 2”). Specifically, for each participant and for each attentional condition, the amplitude of the time-varying signal was averaged across data points within each time window to yield four distinct values per participant (i.e., engage target 1 before switch, disengage target 1 after switch, ignore target 2 before switch, reengage target 2 after switch). These values were compared using a 2×2 repeated measures ANOVA with Bonferroni correction for multiple comparisons.

### Change Point Analysis and Jackknife Procedure

To identify attentional shifts and calculate their latency (relative to the target switch) on both the group and on the individual subject level, we utilized the cumulative sum approach based on Taylor (2000) paired with a jackknifing procedure (Ulrich & Miller, 2001). Using this approach, it is possible to determine whether and if so, when, a change in the data occurred. First, to avoid contamination due to RDK onset, offset and initial attentional engagement, the first and last 0.5 seconds of each trial were excluded from the analysis (Vieweg & Müller, 2021). Then the difference between the averaged signal across time points of each attentional condition and between each point along the signal were computed. Next, a cumulative sum was calculated such that these differences were added consecutively. Finally, a magnitude of change estimator was established by calculating the difference between the highest and lowest values along the cumulative sum curve. For a change point to be considered **valid**, we applied specific criteria based on our normalization procedure. Since only a value > 1 indicates attentional gain and only a value < 1 indicated attentional suppression, valid disengagement was only considered if more than 50% of the data points **before** the change were **above** 1 and more than 50% of the data points **after** the change were **below** 1.

Similarly, valid reengagement was defined by a switch from more than 50% of points **below** 1 before the change, to more than 50% of points **above** 1 after the change. Moreover, a minimum threshold of 2.5 seconds post-RDK onset was set for change point detection. This threshold reflects the task design, where the cue change occurred at a mean of 3 seconds (±0.5 s jitter). If these criteria were not met, the change point was disregarded. The significance of change was determined by means of bootstrap analysis with 10,000 iterations such that the 2560 data points for each attentional condition were randomly reordered. The same cumulative sum procedure was performed for each iteration and a magnitude of change estimator was calculated. To determine whether a significant change has occurred, the number of estimators in the randomized data smaller than the magnitude of change estimator in the original data was divided by the number of bootstraps (10,000) and multiplied by 100 to produce a percentage of estimators smaller than the real estimator. If a change point exceeded a 99% confidence threshold, it was considered significant, and its timing was extracted for further analysis. This reflects the confidence level that a real change has occurred. If a significant change was indeed observed, the maximum point in the cumulative sum curve was extracted and used as an estimate of when this change has occurred. Since this procedure was applied separately for each attentional condition, it allowed us to determine whether attentional disengagement (i.e., shifting attention away from target attended before the switch), reengagement (i.e., shifting attention towards a target that was unattended before the switch), or both occurred and if so, when they occurred along the experimental trial timeline.

To ensure the reliability and consistency of the identified change points at the group level, we employed a **jackknife procedure** (Ulrich & Miller, 2001). This approach involved systematically leaving out one participant at a time, recalculating the group averages, and estimating change points for each iteration. This provided a distribution of group-level change points, from which we calculated mean change points and 95% confidence intervals (CI) across participants. The consistency of the change point estimates was assessed based on the variability in these jackknife iterations, allowing us to verify the robustness of our findings across the sample.

### Signal-to-Noise Ratio Analysis

To ensure that inter individual differences in attentional patterns were not attributable to individual variations in Signal-to-Noise Ratio (SNR), we calculated SNR values for each participant by extracting the power in the stimulation frequency (8, 10, 12, 15 Hz; i.e., the signal) and dividing it by the average power of neighboring frequencies (±1 Hz; i.e., the noise). The power values were extracted as described in section SsVEP signal quality evaluation. The SNR values obtained for each stimulation frequency were then averaged to obtain a single SNR score per participant. Subsequently, a repeated measures ANOVA was employed to examine the SNR scores in relation to the four distinct patterns of attentional disengagement and reengagement identified (see Results section Change point analysis and jackknife procedure).

## Results

### Behavioral Accuracy

Participants were successful at performing the visual task, as indicated by their high behavioral accuracy scores (M = 0.85, SD = 0.13, range: 0.44 – 0.98). The data from one participant who performed poorly on the task (accuracy score = 0.44) was excluded from further analysis leading to mean accuracy scores of M = 0.86, SD = 0.11, range: 0.64 – 0.98.

### SSVEP Signal Quality Analysis

The steady-state visual evoked signal is illustrated in Figure 3a. Reliable peaks at the four stimulation frequencies (8, 10 12 and 15 Hz) are clearly seen in the FFT power spectrum (Figure 3b). Figure 3c shows the topography of activity in each stimulation frequency.

**Figure 3.**
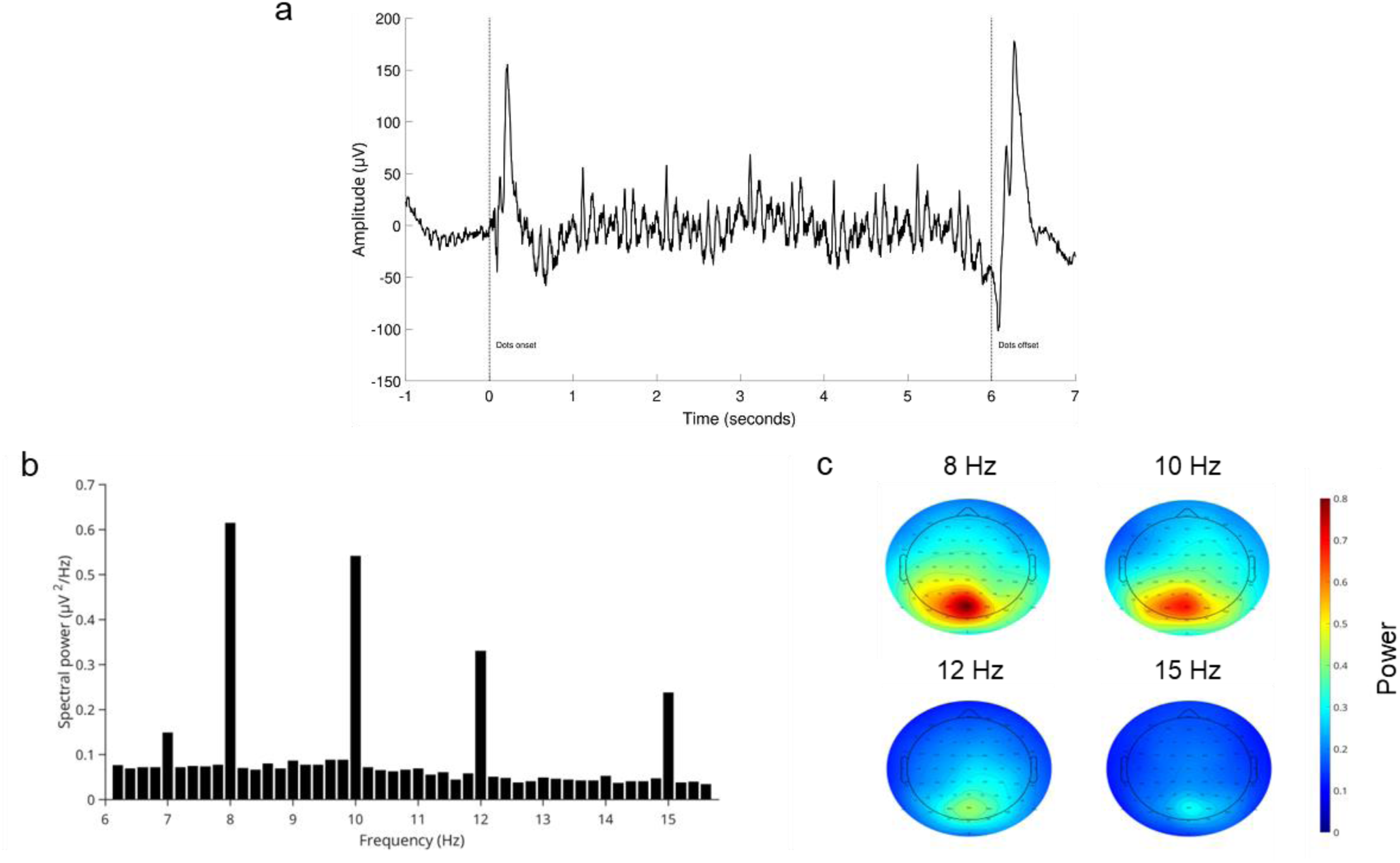
Illustration of the steady-state visual evoked signal. (a) Time domain data from posterior electrodes (’O1’, ’Oz’, ’POz’, ’O2’) averaged across trials, attention conditions and across subjects and the FFT power spectrum (b). Reliable peaks in the power spectrum are evident at the stimulation frequencies (8, 10 12 and 15 Hz) (c) Topographic plots of the averaged power (across trials, conditions and subjects) in each stimulation frequency.

### Time Course and Temporal Dynamics of Attentional Shifts

The final number of trials did not differ significantly between attentional conditions (“engage target 1 - disengage target 1”; “ignore target 2 - reengage target 2”); mean number of first target trials = 79.62, SD = 6; mean number of second target trials = 80.41, SD = 6.17, p > .05, paired t-test).

The time courses of the group-averaged signal along the experimental trial for each attentional condition are presented in Figure 4a. These results align with our hypotheses and provide a detailed picture of the temporal dynamics of attentional engagement, disengagement, suppression, and reengagement. For statistical analysis, the activity from each attentional condition was averaged across time points within the two time windows: 2 s before the target switch and 2 s after the target switch (see section Statistical Analysis), resulting in a 2×2 within-subject repeated measures design, of two time windows (before switch, after switch) by two attentional conditions (Figure 4b). A significant interaction effect was found between time window and attentional condition *F*(1, 28) = 29.31, p < .001, partial η² =.511.

**Figure 4.**
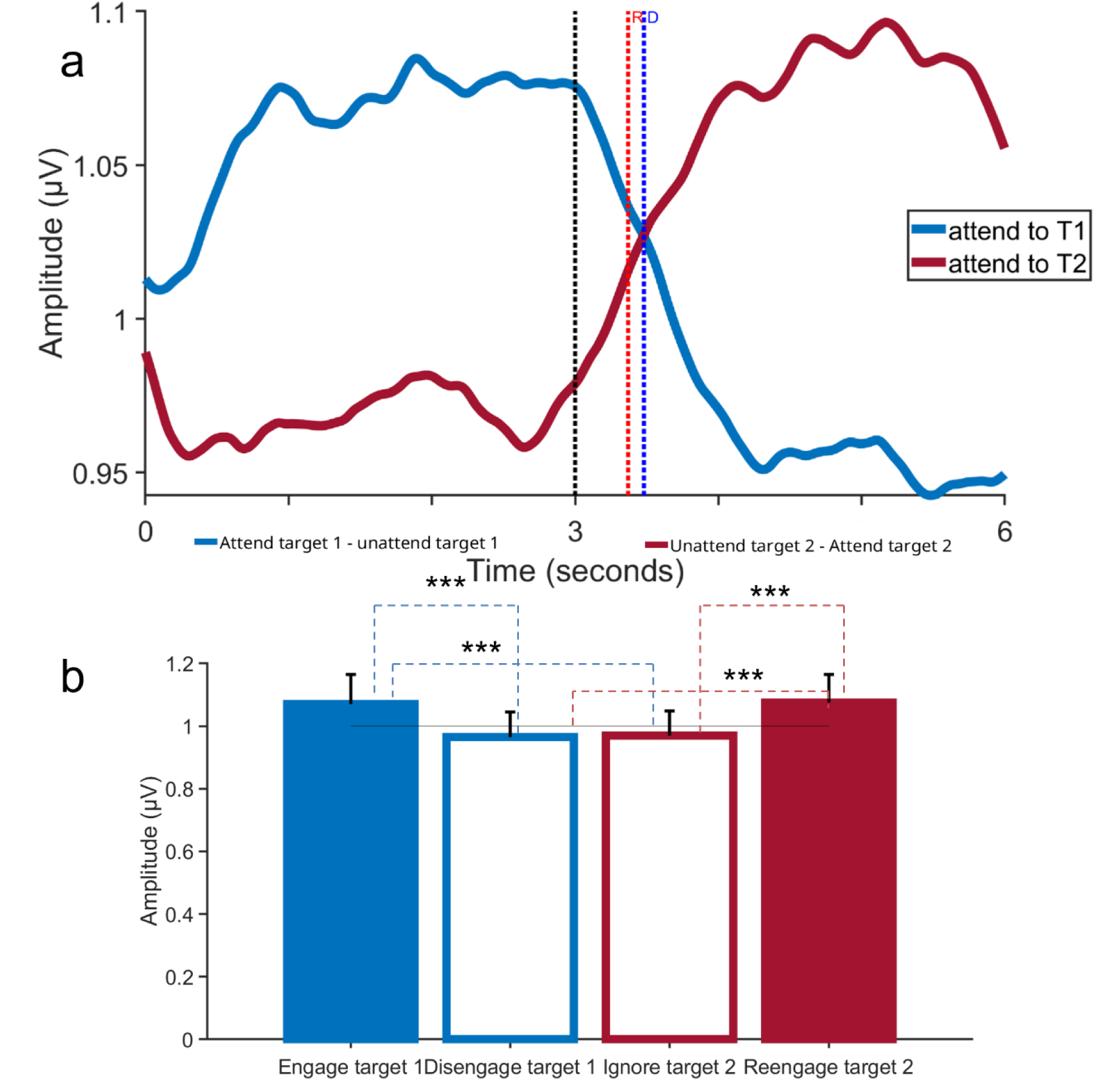
Time course of attentional engagement, disengagement and reengagement. (a) Temporal dynamics of the ssVEP response (Hilbert envelope) in the two attentional conditions across the experimental trial. The blue line represents the first attentional condition (“Attend target 1 - Unattend target 1”) and the red the second attentional condition (“Ignore target 2 - Attend target 2”). In the first time window (0–3 s), **engagement** with target 1 (blue line) is reflected by an increased response, while the suppressed response to target 2 (red line) remains low. Following the target switch (black dashed line at ∼3 s), **disengagement** from target 1 is observed as a decline in the response (blue), while **reengagement** with target 2 is marked by a subsequent increase in the response (red). Significant change points for disengagement (D) and reengagement (R) are indicated by the blue and red dashed lines, respectively. (b) Group averaged activity in each attentional condition in the 2 s time windows for the two attentional conditions. In the first time-window, **engagement** with target 1 is the significantly higher response to target 1 (blue full bar) compared to the suppressed response to target 2 (red empty bar). Following the switch, reengagement with target 2 is reflected in its significantly higher response (red full bar) compared to its earlier suppressed state (red empty bar), while disengagement from target 1 is evident in its reduced response (blue empty bar) compared to its earlier engaged state (blue full bar); ***p < .001.

Pairwise comparisons using Bonferroni corrections revealed significant differences between attentional conditions within each time window indicating engagement and suppression. In the first time-window, the response to target 1 (“engage target 1”; Figure 4b, blue full bar) was significantly higher than the response to target 2 (“ignore target 2”; Figure 4b, red empty bar; p < .001), indicating that target 1 was engaged while target 2 was suppressed. In the second time-window, following the switch, the response to target 2 (“reengage target 2”; Figure 4b, red full bar) was significantly higher than the disengaged response to target 1 (“disengage target 1”; Figure 4b, blue empty bar; p = .002), reflecting the reallocation of attentional resources.

To assess disengagement, pairwise comparisons using Bonferroni corrections tested the difference within the first attentional condition (“engage target 1 - disengage target 1”) between the two time-windows with significant differences observed. Specifically, the magnitude of response to target 1 before the switch (“engage target 1”; Figure 4b, blue full bar) was significantly higher than the response after the switch (“disengage target 1”; Figure 4b, blue empty bar; p < .001). This decrease in response highlights the process of disengagement, where attention is withdrawn from the first target.

To assess reengagement, pairwise comparisons using Bonferroni corrections tested the difference within the second attentional condition (“ignore target 2 - reengage target 2”) between the two time-windows with significant differences observed. A significant increase in response to target 2 was observed after the switch compared to before the switch (Figure 4b, red empty bar vs. red full bar; p < .001). This increase marks the reallocation of attentional resources to the new target, reflecting the process of reengagement.

To test the occurrence and temporal dynamics of attentional shifts, each attentional condition was subjected to the cumulative-sum change-point analysis (as detailed in the method section in Change point analysis and jackknife procedure). This analysis allowed us to identify significant time points of attentional disengagement and reengagement, both relative to the trial start and the mean switch point (set at 3000 ms). This was done on the group level, as well as on the individual level.

For the group-level data, significant attentional shifts were observed. When measured relative to the start of the trial, attentional disengagement occurred at 3,480 ms, while reengagement occurred at 3,370 ms. Relative to the mean switch point (3,000 ms), disengagement occurred 480 ms after the switch, and reengagement occurred 110 ms earlier (i.e., 370 ms after the switch - see Figure 4a, blue and red dashed lines).

At the individual level, the **mean change points** for disengagement occurred at **3,510 ms** (SD = 173.24) relative to the trial start, and reengagement occurred at **3,375 ms** (SD = 237.71). When calculated relative to the mean switch point, disengagement occurred **510 ms after the switch** (SD = 173.24), while reengagement occurred **375 ms after the switch** (SD = 237.71). The highly close adherence to the group level mean reflects the consistency of the overall temporal dynamics across levels of analysis.

The results obtained from the jackknife procedure further confirmed the robustness of these findings, providing confidence intervals to assess the stability of the group-level results.

Specifically, the jackknifed data revealed that disengagement occurred at 3,480 ms (95% CI: 3,410 - 3,540 ms) and reengagement occurred at 3,370 ms (95% CI: 3,320 - 3,410 ms) when measured from the start of the trial. Relative to the mean switch point (3,000 ms), disengagement occurred 480 ms after the switch (95% CI: 410 - 540 ms), while reengagement occurred 370 ms after the switch (95% CI: 320 - 410 ms). The close alignment between the jackknifed results and both group and individual data highlights the reliability and precision of the identified attentional shift latencies. The jackknife procedure, providing confidence intervals, offers additional support for the consistency of these time points across subsamples.

On the individual subject level, our analysis revealed four possible patterns of attentional shifts among participants (Table 1). In 41.38% of participants both disengagement and reengagement occurred; 13.79% of the participants displayed only disengagement; 10.34% displayed only reengagement and in 34.48% neither shifts were displayed.

A closer investigation of the participants who successfully performed both attentional disengagement and reengagement revealed an additional distinction (Figure 5): Specifically, **50% of participants** (n = 6) exhibited reengagement occurring **after** disengagement (positive Δ values), while the remaining **50% of participants** (n = 6) demonstrated reengagement occurring **before** disengagement (negative Δ values). The Δ difference was calculated as **reengagement - disengagement**, with values ranging from **-917.97 ms to 937.50 ms**. The mean Δ for participants who exhibited both shifts was **-90.49 ms (SD = 346.53)**, closely aligning with the group mean of **-110 ms**. This range reflects substantial individual variability in the timing and sequencing of attentional shifts.

**Figure 5:**
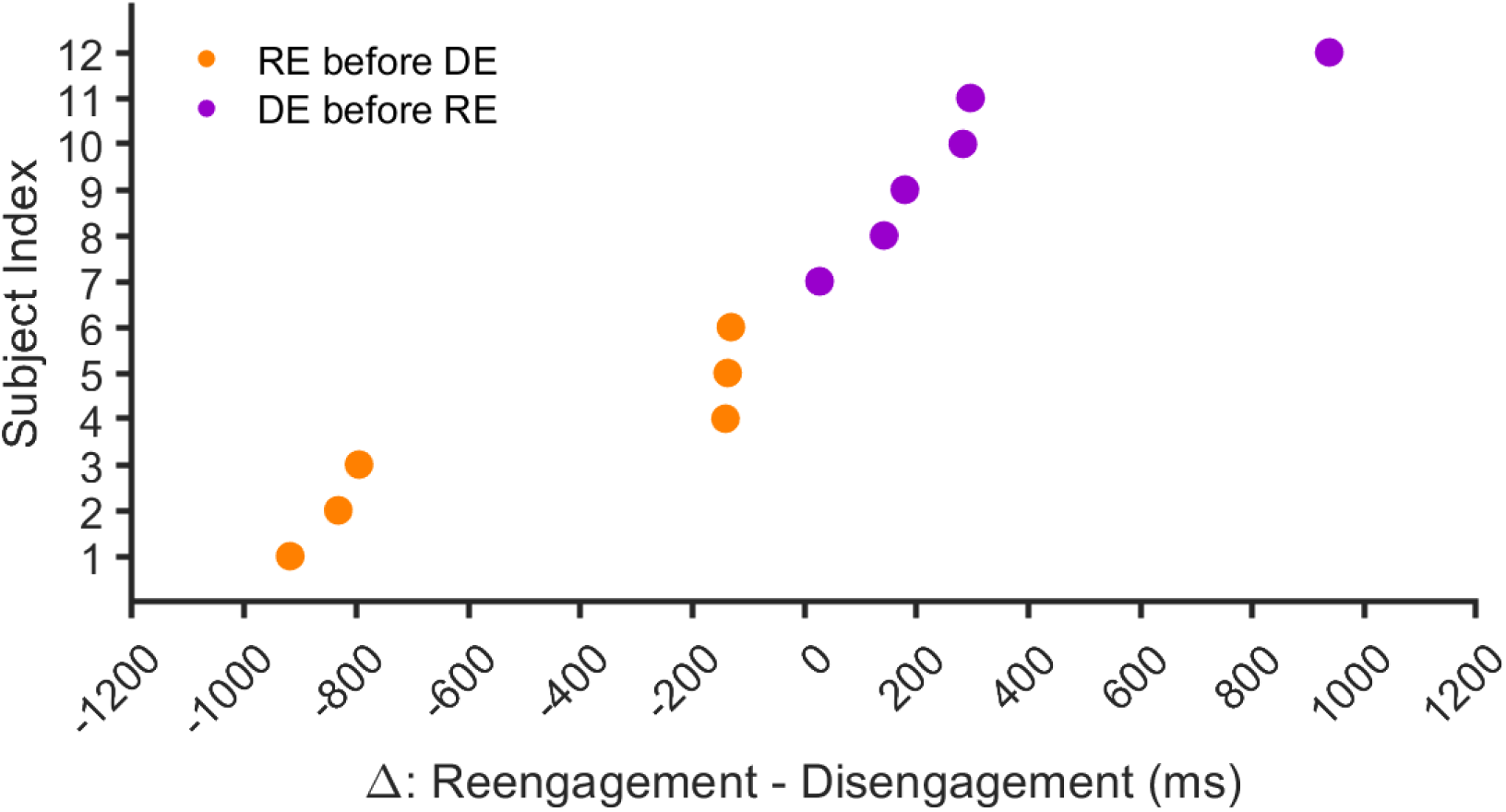
Significant change points for subjects successful in both attentional disengagement and reengagement. **Δ** values (Reengagement - Disengagement) are shown for individual participants. Positive **Δ** values (purple) indicate disengagement (DE) occurred before reengagement (RE), while negative **Δ** values (orange) indicate reengagement preceded disengagement.

No significant differences in SNR (section Signal-to-Noise ratio analysis) were observed among the four emerging attentional shifts patterns described above p > 0.5. This confirms that individual differences in attentional shift patterns are not attributable to variability in signal quality, further supporting the robustness of the identified patterns.

## Discussion

The main goal of the present study was to measure and discern the processes of attentional engagement, disengagement, and reengagement, as well as their corresponding latencies, at both the group and individual level, utilizing the excellent SNR and temporal precision of ssVEPs. To this end, we utilized an integrated approach combining well-validated methods in an experimental framework that enabled us to characterize and quantify the time course of attentional shifts. As hypothesized, on the group level, we initially observed increased ssVEP response to an attended stimulus (attentional engagement), which subsequently declined when the focus was shifted away from it (attentional disengagement). In a similar vein, when a second stimulus became the new target of attentional focus (following a target switch), we observed an increase in response compared to the one measured before it became the attended target (attentional reengagement).

In alignment with previous studies (e.g., Müller et al., 2016; Vieweg & Müller, 2021), on the group level analysis, attentional reengagement temporally preceded disengagement, suggesting that the process of shifting attention to a new target begins before fully disengaging from the previous one. This temporal order reflects the independence of attentional suppression and facilitation processes, as reengaging with a new stimulus does not necessitate the completion of disengagement from a previous one (Andersen & Müller, 2010).

Interestingly, individual-level analysis revealed diverse and nuanced patterns in attentional shifts. While some individuals showed the group level pattern, with reengagement occurring before disengagement, in others only a single shift (disengagement or reengagement), or neither shift was observed. Furthermore, the timing and order of these shifts relative to the target switch varied widely among individuals. Within participants that completed both shifts successfully, half reflected the order of the group data (reengaging prior to disengaging) and the other half reflected the opposite order. Moreover, the latency difference between the shifts among those who completed both shifts was highly variant (spanning between -917.97 ms to 937.50 ms). These substantial profiles, latency and order differences underscores the variability in the temporal dynamics of attentional shifts, suggesting that distinct attentional strategies or cognitive demands may underlie these patterns. By capturing such fine-grained differences at the individual level, our approach provides a valuable framework for exploring the temporal dynamics of attentional shifts at both the group and individual level.

Such variability is particularly relevant when considering the role of attentional shifts in adaptive and maladaptive psychological functioning. Efficient attentional control is a central pillar to the ability to navigate successfully in complex environments, and as previously mentioned disruptions in these processes are frequently linked to different psychopathological conditions (e.g., Abado et al., 2020; Amir et al., 2016; Bar-Haim et al., 2007; De Lissnyder et al., 2011). On the other hand, different interventions like mindfulness-based interventions or attention bias modification aim to enhance the flexibility and efficiency of attentional shifts, emphasizing their role in fostering psychological resilience and well-being (Dahl et al., 2020; Lutz et al., 2015).

Importantly, while group-level findings provide valuable insights into general patterns, they may obscure meaningful individual differences that are critical for understanding how attentional shifts operate across diverse populations. Therefore, as highlighted in recent literature, developing tools that reliably capture individual-level outcomes is essential for evaluating the effectiveness of interventions such as those discussed above (Davidson & Kaszniak, 2015; MacLeod et al., 2019). By bridging group-level findings with individual-level patterns, the methods developed in this study can highlight practical pathways for developing interventions that target attentional control.

Our methodological approach has demonstrated the potential for capturing both group and individual differences in attentional shifts. However, in order to determine whether this approach can provide a reliable measure of attentional shifts both in research and clinical applications, future studies should focus on testing its’ effectiveness in various contexts. For example, utilizing it along with psychological assessments as well as testing its’ sensitivity and reliability across different attentional control interventions (such as the ones described above). Notably, the need for an integrated approach assessing attentional shifts not only at the behavioral level, where cognitive effort can often be masked, but also at the neural activity level where the underlying processes can be directly observed, has been frequently addressed (Eysenck et al., 2022; Lutz et al., 2015; Vago et al., 2019).

Several limitations should be taken into consideration when interpreting the results of our study. First, our experimental design may have introduced a bias in favor of the first attentional target, since the first cue arrow presented before the RDK onset was always identical to the one denoting the first target. While our analysis shows that the overall responses to target 1 and target 2 did not differ significantly (Figure 4b), future designs should account for such a potential bias. Second, in the current experimental design, attentional shifts were possible between different features (two colors – red and green) in the same and / or in different location (left or right side of a central fixation point). Except for the constraint that a target cannot remain the same before and after the attentional switch, all shifts were possible. While this design enabled us to quantify the processes we intended to examine (namely attentional engagement, disengagement, and reengagement), there are nuances regarding specific patterns of attentional shifts in the different attention allocation options introduced by our design, that require higher statistical power (i.e., more trials of each type of shift) in order to be profoundly examined. For example, in Adamian et al., 2019, dividing attention between features did not result in a similar outcome as dividing attention between locations. Therefore, it is possible, for example, that in our study participants displaying a pattern of reengagement alone (i.e., no disengagement) may have actually divided their attention without cost, i.e., they widened their attention focus to include the new target, rather than switched it between the previous and the new target. Thus, further studies with a larger number of trials for each type of shift, whether feature-based or spatial, are needed in order to characterize attentional processes more fully.

In conclusion, this study represents a step forward in understanding temporal dynamics of attentional shifts. Our findings highlight the importance of examining attentional shifts on both the group and individual levels, while demonstrating the independent and gradual nature of these processes. This approach may not only provide significant insights when assessing individual differences, but can also be valuable in improving the precision of interventions aimed at enhancing attentional control, with implications for both research and clinical practice.

## Author Contribution

Moran Eidelman-Rothman: Conceptualization, Methodology, Software, Formal analysis, Investigation, Data curation, Writing – original draft, Writing – review & editing, Visualization, Project administration. Omer Reuveni: Conceptualization, Methodology, Software, Formal analysis, Investigation, Data curation, Writing – original draft, Writing – review & editing, Visualization, Project administration. Andreas Keil: Conceptualization, Methodology, Formal analysis, Validation, Software, Resources, Writing – review & editing. Lior Kritzman: Conceptualization, Methodology, Software, Formal analysis, Investigation, Data curation, Writing – review & editing, Visualization. Dominik Freche: Conceptualization, Methodology, Investigation, Data curation, Software. Hadas Okon-Singer: Conceptualization, Resources, Writing – review & editing, Supervision. Nava Levit-Binnun: Conceptualization, Methodology, Resources, Writing – original draft, Writing – review & editing, Supervision, Funding acquisition.

## Acknowledgments

This study was supported by the Joy Academic Grant Program and the Sagol family foundation. We extend our gratitude to Amotz Taub-Tabib for his invaluable insights and dedication. Special thanks to our research assistants Adi Gnapp, Eden Tzadik and Galit Pesyachov, for their exceptional contributions and commitment to this study.

