## Supplemental Figure 1 for "Quantifying Variability in Attentional Shifts During Target Switching Using Steady-State Visual Evoked Potentials"

### Supplementary

#### Individual level analysis: Change point analysis using cumulative sum.

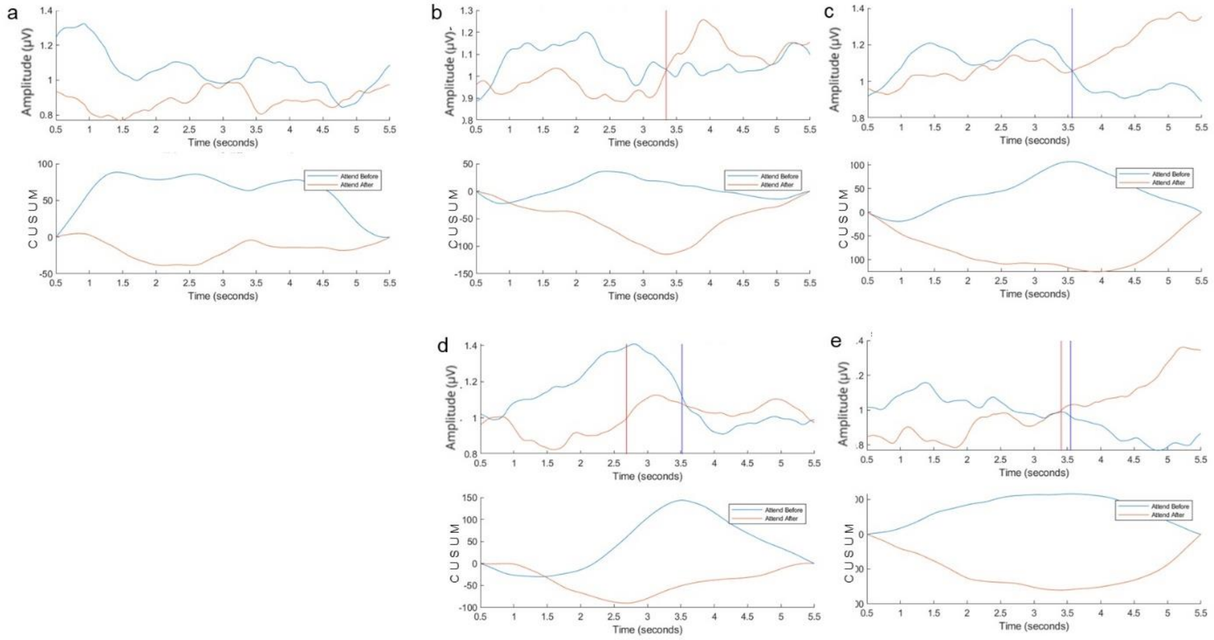


**Figure 1.** Examples of individual patterns of significant change points. In each example, the top panel presents the the ssVEP downsampled response (Hilbert envelop) in the two attentional conditions. The bottom panel represents the cumulative sum (CUSUM) for the two attentional conditions (see section Change point analysis using cumulative sum). Blue represents attentional condition 1 and orange represents attentional condition 2. The significant change point along the response is represented by vertical lines, (a) Participants with no significant shifts (b) participant exhibiting disengagement alone (change point marked with red vertical line) (c) participant exhibiting reengagement alone (change point marked with blue vertical line) (d) and (e) participants exhibiting both disengagement and reengagement. The two examples showed in d end e demonstrate that even when exhibiting both disengagement and reengagement, these can occur at different latencies and in different order.
